# Taxonomically mixed blue mussel *Mytilus* populations are spatially heterogeneous and temporally unstable in the subarctic Barents Sea

**DOI:** 10.1101/2022.12.08.519596

**Authors:** Julia Marchenko, Vadim Khaitov, Marina Katolikova, Marat Sabirov, Sergey Malavenda, Michael Gantsevich, Larisa Basova, Evgeny Genelt-Yanovsky, Petr Strelkov

## Abstract

Subarctic populations of blue mussels represented by “cryptic” species *Mytilus edulis* (*ME*) and *M. trossulus* (*MT*) have been studied less intensively than Arctic and boreal populations. Ecological features of *ME* and *MT* in sympatry are poorly known everywhere. The knowledge about mussels at the northeasternmost boundary of the Atlantic littoral communities on Murman coast of the Barents Sea is based on data obtained 50-100 years ago. Our study provides the first insight into the long-term dynamics of the Barents Sea mussels, the habitat segregation of *ME* and *MT*, and the interannual dynamics of their mixed settlements. The Tyuva Inlet (Kola Bay), which is 3 km long, was used as the study site. Mussels were found everywhere in the littoral and the sublittoral down to a depth of 4 m. Their characteristic habitats were sandbanks, littoral rocks, sublittoral kelp forests and “the habitat of the mussel bed” in the freshened top of the inlet. The main spatial gradients explaining the variability of demographics of the settlements (abundance, age structure, size) were associated with the depth and the distance from the inlet top. *ME* and *MT* were partially segregated by depth: *ME* dominated in the sublittoral and *MT*, in the littoral. In addition, *ME* dominated both in the littoral and in the sublittoral parts of the mussel bed. The ratio of species in the mixed settlements varied over time: between 2004 and 2010 the proportions of *MT* decreased everywhere, by 22 % on average. Historical data indicate that the abundance of the Murman mussels declined sharply between the 1960s and the 1970s, which coincided with the cooling of the Arctic. It seems that the populations have not completely recovered in the abnormally warm recent decades. The habitat distribution of mussels apparently changed with time, too: unlike today, in the 20th century mussels were rarely observed in kelps. We suggest that the spatial and temporal dynamics of subarctic mussels can be partly explained by the competition between *ME* and *MT* combined with their differing sensitivity to environmental factors.

## Introduction

Populations of blue mussels (*Mytilus* spp.) in the Arctic and in the Antarctic have received much scientific attention in recent decades. Studies have been made in the East Siberian Sea (Gagaev et al., 1994), Northeastern Alaska (Feder et al., 2003), Spitsbergen (Berge et al., 2005; Leopold et al., 2019, Kotwicki et al., 2021), the Pechora Sea (Sukhotin et al., 2008); Northwestern Greenland (Blicher et al., 2013; Thyrring et al., 2015) and, on the other side of the Globe, in the South Shetland Islands (Cárdenas et al., 2020). A keen interest in these populations, most of which had previously been unknown or even non-existent, is due to the fact that they represent the coldest parts of the mussel distribution and are pioneers in the poleward expansion under conditions of the warming climate. Blue mussels from temperate seas have always received much attention, due to their important ecological and economic roles (Gosling, 2021). Significant declines of their populations in some areas such as the Gulf of Maine (Sorte et al., 2017) and the Atlantic coasts of Sweden (Baden et al., 2021) and France (Seuront et al., 2021) have been registered in recent decades and mainly explained by the climate change.

In contrast, recent studies of the subarctic populations of blue mussels are relatively scarce. To note, the Arctic and the Subarctic are defined in this paper according to the Conservation of Arctic Flora and Fauna (https://www.caff.is/). While the subarctic mussels are not entirely neglected, having been studied at the White Sea (Lukanin et al., 2006; Khaitov & Lentsman, 2016), in the northern Gulf of Alaska (Bodkin et al., 2018) and in the Sea of Okhotsk (Selin & Lysenko, 2006; Khalaman et al., 2020), they have clearly been overshadowed by the Arctic and the temperate ones. This is probably due to the facts that the subarctic mussel populations are less interesting biogeographically and less important economically; besides, being long known, they have already been examined at one time or another.

Populations of the Barents Sea coast of the Kola Peninsula, broadly known as the Murman Coast or Murman, at 68-70 degrees north and 31-40 east (**Fig. 1**) are a case in point. The Murman Coast is washed by a warm Atlantic Murman Coastal Current, which is responsible for relatively high sea surface temperatures (SST) for such high latitudes (the long-term SST is +10.2°C for August and +3°C for February in Ekaterininskaya Gavan in the Kola Bay, https://www.seatemperature.org/) and a limited winter ice cover. The Barents Sea is strongly affected by long- and short-term quasi-regular climate fluctuations, with the SST varying by several degrees Celsius on interannual and more than a degree on decadal time scales (Ingvaldsen et al., 2021; Matishov et al., 2012; www.pinro.ru). Murman represents the northeasternmost border of the typical littoral communities of the North Atlantic, with their canopies of fucoid algae, crusts of barnacles and mussels on hard bottoms (Zatsepin et al., 1948; Genelt-Yanovsky et al., 2019 and references therein).

**Figure 1.**
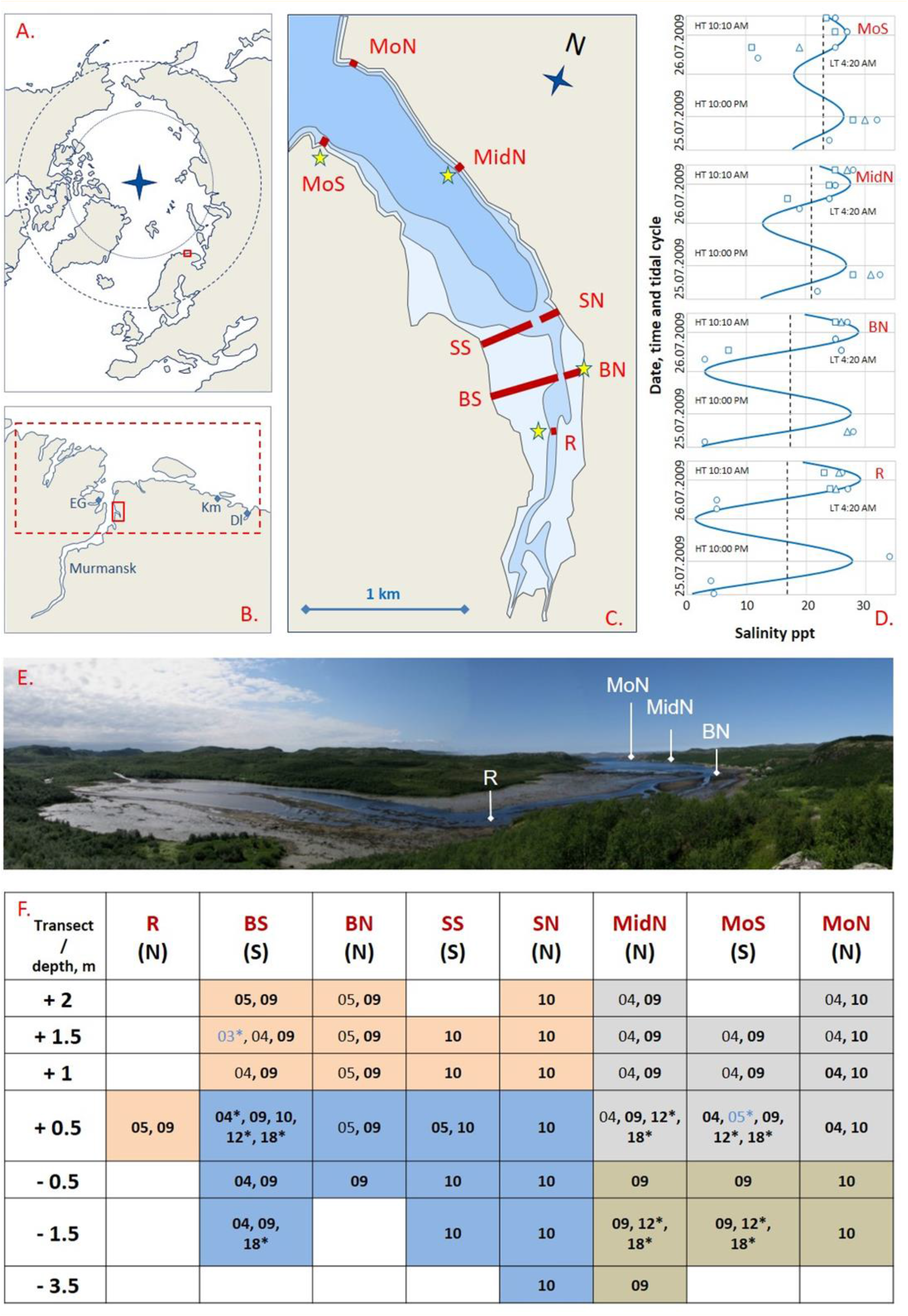
Tyuva Inlet and the mussel sampling scheme. (**a**) Polar view map of the Arctic Ocean. Box indicates the location of Kola Bay. (**b**) Map of the Kola Bay and surroundings. The small box indicates the location of the Tyuva Inlet, and the large one indicates the area for which historical data on mussels were collected. Abbreviated names: EH - Ekaterininskaya Gavan Bight, Km - Klimkovka Inlet, Dl - Dolgaya Inlet. (**c**) Map of the Tyuva Inlet. The littoral, the sublittoral shallower than 10 m and the sublittoral deeper than 10 m are shown in different shades of blue. Red lines labeled by abbreviated names show transects along which mussels were sampled in 2003-2018. Asterisks indicate salinity sampling points. (**d**) Variation of salinity, ppt, in Tyuva surface waters on 25-26 July 2009. Blue curves show predicted salinity, dashed lines – average salinity, signs – empirical data (the shape reflects the depth of sampling: circles – surface, triangles and squares – depth of 1 and 2 m from the surface, respectively) in the four intertidal localities labeled as in **c**. The lowest (LT) and highest (HT) tidal times are indicated. (e) Top of the Tyuva Inlet by low water on 21.07.2009. Intertidal flats are visible. Position of some transects is shown. For scale, the distance between R and BN is 250 m, between MidN and MoN is 950 m. (**f**) Locations and years of mussel surveys in the Tyuva Inlet in 2003-2018. Columns are transects, notations as in **c**, N and S denote the northern and the southern coast, respectively. The lines are depth horizons from the chart datum (negative values denote sublittoral position). Numbers are years of studies (03 – 2003, 04 – 2004, etc.). Blue font and (or) asterisks indicate years of sampling for genetics, black font, for demography, and bold font, for taxonomic structure by morphotypes. Cell filling reflects mussel habitat. Pink – littoral sandbanks, orange – sublittoral kelp forests, gray – rocky littoral, blue – habitat of the mussel bed.

There are abundant data on coastal macrobiotic organisms of Murman, resulting from extensive phenomenological studies conducted there in the early 20th century, mostly at the former Murman Biological Station in Ekaterininskaya Gavan (Guryanova et al., 1928, 1929; Zatsepin et al. 1948; Matveeva, 1948; note that the publications of 1948 are based on the pre-World War II data). According to these studies, mussels were among the most conspicuous littoral species. At the same time, they were rare in the sublittoral, except in mussel beds in the river mouths (Guryanova et al., 1926; Matveeva, 1948). After that, the Murman mussels were considered in a few studies, whose main conclusion was that mussel abundances decreased dramatically between 1960s and 1970s along the entire coast and did not recover until 1980s (Antipova et al., 1984) or even late 1990^th^ (Strelkov et al., 2001). This decrease was assumed to be related to a prolonged period of low sea water temperatures that has started in 1960 and was, supposedly, unfavorable to mussels (Antipova et al., 1984).

To sum up, the knowledge of Murman mussel ecology is mainly based on the data that are 50-100 years old. How well it reflects the current situation is anybody’s guess. In our opinion, there are at least three reasons to believe that this knowledge is outdated: recent environmental changes, modifications of the sampling methods and taxonomic changes. To begin with, the first two decades of the 21^st^ century were unprecedentedly warm in the Barents Sea (Marshall et al., 2016; Ingvaldsen et al., 2021; www.pinro.ru). If mussel abundance in the Barents Sea indeed positively depends on temperature (Antipova, 1984), we may expect a recovery after the supposed population decline in the 1960^th^ and a high mussel abundance. Secondly, in earlier studies sublittoral mussels were sampled by dredges, while today they are usually picked by divers. These differences probably impact the inferences from the studies.

Last but not least, the knowledge of mussel ecology dating back to the 20^th^ century is likely to be flawed because of the upheavals in mussel taxonomy that have occurred since that time. In the late 1980s, the Arctic-boreal mussel species *Mytilus edulis* was divided into *M. edulis* (hereinafter, *ME*) and *M. trossulus* (*MT*) based on genetic data (McDonald et al., 1991). In origin, *ME* and *MT* are vicarious species that have been evolving independently since the Pliocene in the Atlantic and the Pacific Ocean, respectively. It was only postglacially that *MT* invaded the Atlantic sector (Laakkonen et al., 2021 and references therein). Today, *ME* and *MT* co-occur and hybridize in many areas of the Atlantic and the neighboring Arctic, including Murman (Väinölä & Strelkov, 2011; Wenne et al., 2020). *MT* is less thermophilic: it does not spread as far south as *ME* in the Atlantic (Wenne et al., 2020 and references therein) and shows a poorer physiological performance at elevated temperatures (Rayssac et al., 2010; Fly & Hilbish, 2013; Bakhmet et al., 2022). Otherwise, *ME* and *MT* appear to play similar ecological roles in their native oceans, and their ecological differences can only be assessed in sympatry (Riginos & Cunningham, 2005).

Theoretically, some ecological differences between *ME* and *MT* should be expected, since they must inevitably be competing for resources. However, little is known on this topic in general (see Riginos & Cunningham, 2005; Katolikova et al., 2016 for review), and nothing at all in the Barents Sea. This lack of knowledge is partly due to the fact that *ME* and *MT* are “cryptic” species, with no diagnostic morphological features, and the genotyping methods traditionally used for their identification are very laborious (Katolikova et al., 2016). It has recently been shown that *ME* and *MT* in the Murman populations differ by frequencies of shell morphotypes defined as absence or presence of an uninterrupted prismatic strip under the ligament on the inner side of the shell. In brackish localities (<30 ppt) the differences approach 65%, while in saline localities they make up only 18% (Khaitov et al., 2021). This means that in saline localities individual mussel assignment to one of the two species based on morphotypes is ineffective; however, the proportions of the species in samples can be predicted based on the morphotype frequencies fairly accurately (Khaitov et al., 2021).

Here we present the results of a new phenomenological study of Murman mussels, which was driven by two compelling gaps: the lack of up-to-date data on mussels from this area and the scarcity of information about habitat preferences of *ME* and *MT* in sympatry. The Tyuva Inlet in the northeastern part of Kola Bay (**Fig. 1a, b**) was chosen as a study site for the following reasons. (1) Morphologically and oceanographically, the Tyuva is a typical Murman small inlet, with a deep rocky entrance and a shallow sandy top freshened by the inflowing river (Derjugin, 1915). (2) It is one of the few relatively undisturbed inlets in the Kola Bay where research is still possible. In comparison, the Ekaterininskaya Gavan (8 km from the Tyuva directly across the Kola Bay, **Fig. 1b**), where a biological station was formerly situated, has become a paramilitary zone and is inaccessible. (3) Both *ME* and *MT* were recorded in the Tyuva by the geneticists (Väinölä & Strelkov, 2011), which makes the inlet a suitable place for a study of sympatric mussels. (4) In retrospect, our interest in Tyuva was to a great extent due to reports of local residents about a large mussel bed there, supposedly the largest in the entire Kola Bay. Using abundant material from the Tyuva (259 quantitative samples from 43 mussel settlements) accumulated in 2004-2018, we could describe the relationships between the taxonomic structure and the basic demographic characteristics (abundance, age structure, size at certain age) of the settlements on the one hand and the key environmental factors, as well as time, on the other hand. We also compared the patterns revealed in our study with those described in the past, for which purpose we summarized the old data on regional mussel populations.

## Materials and Methods

### Tyuva Inlet

The Tyuva Inlet is 3 km long and 0.7 km wide at its widest. The shores of the outer part of the inlet are steep, the littoral zone is narrow, the dominant bottoms are formed by pebbles and rocks. Fucoid algae are abundant on the littoral. Towards the top of the inlet, the shores become more gentle. The inner part of the inlet is shallow, with unconsolidated bottoms and broad sandbanks up to 450 m wide. Fucoids are more scarce there. The river (the common bed of the Bolshaya Tyuva and the Malaya Tyuva that join at the very top of the inlet), with an annual runoff of about 0.7 km^3^, flows across the shoals. The tidal amplitude in this area of the Kola Bay is 1.1-3.7 m, surface salinity at a distance from the river mouths is 31-32 ppt, ice conditions are fast ice in cold winters (Morozov, 1901; Derjugin, 1915; Guryanova et al., 1929; Mytaev, 2014; Shavykin, 2018; **Fig. 1c, e**).

### Mussel sampling and processing

Mussels were studied in the Tyuva Inlet in six different years: 2004, 2005, 2009, 2010 (in July), 2012 (in September), and 2018 (in July). The samples were collected at a low tide to accurately predict the depth based on the tidal data for the Ekaterininskaya Gavan. Quantitative samples, 1-18 per sampling locality, were taken randomly by a core of 0.01-0.25 m^-2^. Qualitative samples for genetics were taken at the chosen localities in different years. A complete description of the sampling design is presented in **Fig 1f** and **Stable 1**. The community of mussels inhabiting a particular locality is referred to in the text as a “settlement”.

In 2004-2005 distribution of mussels was mapped: i) by on-shore observations at the tidal zone along the entire inlet; ii) by SCUBA divers in the subtidal zone in the inner part of the inlet where a large littoral-sublittoral mussel bed (hereinafter, the Bed) was located. Twenty-three mussel settlements were sampled from different depths at three visually identified characteristic habitats: rocky littoral, littoral sandbanks and the “habitat of the Bed”, defined as a broad area of the seabed with a dense settlement of predominantly large mussels, enriched with black sulfuric silt usually accumulated in mussel beds (**STable 1**; photos of habitats are provided in the **SFig. 1**).

In 2009-2010, in addition to the three types of habitats defined during the previous period, mussels settlements in subtidal kelp forests represented by *Alaria esculenta, Saccharina latissima* and *Laminaria digitata* were found by SCUBA diving at 0-4 m depth in the outer part of the inlet. Sampling was performed along seven transects oriented perpendicularly to the coastline at seven depths: +2, +1.5, +1, +0.5, −0.5, − 1, −1.5 m (in relation to chart datum, negative values denote sublittoral position; the range was chosen to cover all the depths inhabited by mussels). Mussels were also sampled from the last upriver littoral settlement, designated as R+05 (**Fig. 1c**,**f**). The choice of transects and the sampling depth reflected the need i) to replicate the sampling design from 2004-2005 and ii) to account for the diversity of mussel habitats. Totally 43 settlements were sampled (**Fig. 1f, STable 1**).

In 2012 and 2018 five and six settlements were resampled, respectively. In 2018 additional qualitative samples were obtained from the same settlements for genetic analysis (**Fig. 1f, STable 1**).

Mussels from each sample were counted and weighted. The maximal anterior-to-posterior length of each mussel (hereinafter, “shell length”) was measured using calipers or dissecting microscope micrometer with a precision up to 0.1 mm. Age of mussels was assessed by counting “winter rings”: marks of winter growth delays on shells as in Sukhotin et al., 2007. For mussels aged 4-7 years the shell morphotypes (E-morphotype, more characteristic of *ME*, or T-morphotype, more characteristic of *MT*) were determined as in Khaitov et al., 2021. Only medium-aged mussels were used in the taxonomic analysis in order to avoid the bias due to a possible association between size and morphotypes in conspecific mussels (see Khaitov et al., 2021 for details).

### Environmental parameters assessment

Every mussel settlement was characterized by the following environmental parameters: *Depth* – height/depth from chart datum, m; *Bottom* – prevailing ground type (boulders, rocks or sand); *Kelp* – presence/absence of kelps; *Cov* – cover abundance of macrophytes by visual observations rated on a rank scale (1 – <5%, 2 – 5-25%, 3 – 25-50%, 4 – 50-75%, 5 – >75% cover); *Slope* – the degree of bottom slope at the sampling point estimated as the slope value for the tangent at that point on the transect profile. We also considered transect-specific characteristics (the same for all localities along the transect): *Dist* – distance (by the midline of the inlet) between the transect and the settlement closest to the river mouth (R+0.5), m; *Width* – distance from the uppermost to the deepest sampling localities within the same transect, m (roughly proportional to the width of the mussel belt on a given stretch of shore) and *Exp* – the shore exposure (north or south). *Depth, Bottom, Kelp* and *Cov* were assessed in parallel with mussel sampling in 2009-2010. *Dist* and *Width* were taken from the map. *Slope* was calculated from vertical transect profiles reconstructed by depths and geographical coordinates of the sampling localities.

Salinity was monitored throughout 25-26 July 2009 at sampling localities R+0.5, BN+05, MidN+05 and MoS+05 located in different parts of the inlet (**Fig. 1c**). Water samples (52 in total) were taken repeatedly at different phases of the tidal cycle (tide range was 0.2-3.8 m above chart datum), with a bathometer, at the surface and at depths 2 m and 1 m from the surface. Salinity was measured with a refractometer S/Mill-E, Atago, Japan with an accuracy of 1 ppt. To predict salinity at each of four littoral sites, we constructed linear model (see below) in which salinity was treated as the dependent variable and tidal height (*H*) at the time of sampling, according to the tide table for Ekaterininskaya Gavan, and *Dist*, as predictors. After building the model, we predicted salinity throughout the tide cycle for each site.

### Demographic and taxonomic parameters of mussel settlements

The following characteristics of mussel settlements were considered: *B* – biomass, g*m^-2^, *N* – total density, ind*m^-2^, *N2-3, N4-6, N7-9, N10* – densities, ind*m^-2^, of mussels 2-3, 4-6, 7-9 and over 9 years old, respectively, *Lma*x – the length of the largest mussel, *L5* – mean length of mussel at the age of five years (the oldest age class present in most of the samples), *GI* – the mussel size at age index, *Ptros* – proportion of *M. trossulus* predicted by frequency of T-morphotypes (see Prediction of taxonomic structure by morphotypes section below). The density of one-year-old mussels was ignored (though it was considered in the calculation of *N*) because of their patchy distribution, which is difficult to account for in a limited sample. *GI*=log (L_∞_ * K), where L_∞_ and K – parameters of the von Bertalanffy equation calculated from the average values of the shell lengths of animals of different ages over 2 years old. *GI* is used here as an indirect measure of mussel growth conditions in settlements. A similar individual-based index known as Overall Growth Performance (OGP) is used in ecophysiology to account for the rate of the organism’s size increase during the lifetime (Brey, 2001 and references therein).

Pooled samples from individual settlements were used for calculations of *Lmax, Ptros* and *GI*. Averaged data on multiple samples from individual settlements were used for calculations of the other characteristics.

### Statistical analyses

All statistical analyses were performed with functions of R statistical programming language (R Core Team, 2020). Multidimensional analyses (CA, CCA, PERMANOVA, SIMPER) were performed by “vegan” package (Oksanen et al., 2020), regression analysis was performed by “glmmTMB” (Brooks et al., 2017) and “mgcv” (Wood, 2011) packages. In all analyses where permutational procedures were implemented, 9999 permutations were set up.

#### Prediction of taxonomic structure by morphotypes

Khaitov et al. 2021 provided formulas to predict the proportion of *MT* (*Ptros*) based on the proportion of T-morphotypes (*PT*) in samples from brackish (<30 ppt) and saline (≥30 ppt) habitats. The salinity boundary between brackish and saline habitats was chosen conventionally, and six samples from the Tyuva used in that study were treated as being from saline habitats. Since mussels in the Tyuva Inlet experience very variable salinity, and the habitat could not be defined as either brackish or saline (**Fig. 1d** and below), we clarified the relationship between *Ptros* and *PT* for local settlements. For this purpose, we have used 15 genotyped samples from the Tyuva Inlet (sample size 30-82 individuals, mean 44, **STable 2**), including nine from the published studies (Bufalova et al., 2005; Väinölä & Strelkov, 2011; Khaitov et al., 2021), stored in collections of the Department of Ichthyology and Hydrobiology (St. Petersburg State University), and six new samples collected in 2018 (**Fig. 1f, STable 1**). New samples were genotyped by the same set of allozyme loci “diagnostic” for the two species (Est-D, Gpi and Pgm) as in the studies listed above. Multilocus genotypes were classified into two categories, those dominated by *ME* genes and those dominated by *MT* genes, using Structure approach (Pritchard et al. 2000) as in Khaitov et al., 2021. For ease of presentation, these categories will be referred to as “*M. trossulus*” (*MT*) and “*M. edulis*” (*ME*) although each could include hybrids in addition to purebreds (Khaitov et al., 2021). To note, hybrids between *ME* and *MT* are rare in the Kola Bay (5-15% by different estimations, Simon et al., 2021; Wenne et al., 2020; Khaitov et al., 2021). The age of mussels was identified and only mussels aged 4-7 years were used in the analysis. Empirical relationships between *PT* and *Ptros* within the three Barents Sea sample sets (15 samples from the Tyuva, 8 samples from saline localities excluding Tyuva and 12 samples from brackish localities from Khaitov et al., 2021) were derived using a regression approach as in Khaitov et al., 2021. In the logistic regression model based on binomial distribution (logit link-function) *Ptros* was considered as a dependent variable, while *PT* and sample set were considered as predictors. Interaction between the predictors was also included in the model.

#### Analysis of population and taxonomic structuring of Tyuva mussels in 2009-2010

To evaluate the population and taxonomic structuring of the Tyuva mussels and to describe how taxonomic structure and demographic characteristics of the settlements were related to the key environmental parameters we used abundant data from 2009-2010. Associations between all demographic, taxonomic and environmental parameters (except salinity) were quantified with the help of canonical correspondence analysis (CCA; Ter Braak & Verdonschot, 1995). Associations between *Ptros* and environmental parameters were also analyzed separately using regression analysis. To compare groups of settlements from different habitats identified visually during sampling (i.e. rocky littoral, sandbanks, kelp forests and the Bed) by demographic parameters and *Ptros*, Permutation Multivariate Analysis of Variance (PERMANOVA; Anderson, 2014) was used.

In the CCA analysis, the matrix of dependent variables contained *Ptros* and all demographic parameters, while the constraints matrix contained all environmental parameters. An optimal CCA model was constructed with the use of forward selection protocol (Blanchet et al., 2008). The statistical significance of the optimal model, individual canonical axes and constraints was assessed by permutation methods (Legendre & Legendre, 2012).

In the regression analysis, a generalized linear mixed model (GLMM) with beta distribution and a logit link-function was used, where *Ptros* was the dependent variable, and environmental parameters were predictors (the values of quantitative environmental parameters were standardized). The transect was included into the model as a random factor influencing the model intercept. Before fitting the model, the set of all predictors was checked for collinearity by calculating the variance inflation factors (*vif*) (Fox 2016). If *vif* exceeded 2 the predictor was excluded. The validity of the final model was inspected by visual analysis of residual plots and the assessment of the presence of overdispersion. Since the test statistic estimated by GLMM corresponds to the Chi-square distribution only approximately (Zuur et al., 2009), we considered p-values less than 0.01 to be significant.

Data preparation for PERMANOVA was as follows. The matrix of dependent variables (the same as in CCA) was transformed (log(x+1)), the Bray-Curtis dissimilarity matrix was calculated and the equality of within-group variance was checked. PERMANOVA was followed by pairwise comparisons of groups. For these multiple comparisons the p-values were adjusted with a Bonferroni correction.

#### Temporal dynamics of the Tyuva mussels in 2004-2018

The choice of strategy for analyzing temporal dynamics was associated with the heterogeneous structure of the data from different study periods. All of the 23 settlements surveyed in 2004-05 were also surveyed in 2009-10, but there were no settlements from kelp forest among them, and only nine settlements were characterized by *Ptros* in 2004-05. Out of the five settlements studied in 2012, all the five were studied in 2009-10 and 2018 but only three were studied in 2004-09. Only three settlements were examined in all the four study periods, among them BS+05 (littoral part of the Bed), which was examined in 2005, 2009, 2010, 2012, 2018 (**Fig. 1f**). In one set of the analyses aimed at identifying trends over the entire observation period, we assumed that the settlements were randomly selected for the study in different years. In another set of analyses, aimed at examining the changes between 2004-05 and 2009-10, we compared only the overlapping sets of the samples.

To assess the variability of mussel demography in the entire material, we applied Correspondence Analysis (CA) based on the matrix of all demographic parameters. The scores of the first CA axis (CA1), which explained the bulk of the total inertia (see Results), were treated as a generalized characteristic of the demographic structure of the settlements. To analyze temporal changes, the scores were used as the dependent variable in the generalized additive regression model (GAM, normal distribution) with Year and *Habitat* as predictors. The smoothers for each habitat were fitted separately and the transect was treated as a random effect (Type I model in Pedersen et al., 2019). Temporal dynamics of *Ptros* was studied separately using a similar approach. The structure of the fitted model was the same but the beta-distribution for the dependent variable was chosen for the GAM construction.

To find out whether the demography of mussel settlements in general and in three different habitats (rocky littoral, sandbanks, the Bed) in particular changed in a unidirectional way between 2004-05 and 2009-10, we analyzed the data on 20 settlements sampled in both periods (3 settlements lacking *GI* and *L5* in 2004 were excluded) using PERMANOVA with two factors, *Habitat* (three levels: rocky littoral, sandbank, bottom) and *Period* (two levels), and the interaction between them. A similarity percentages (SIMPER) analysis (Clarke 1993) was further performed to estimate the contribution of each demographic parameter into the formation of differences between the two temporal periods. Data preparation and assumption testing for PERMANOVA and SIMPER were identical and the same as described above. We also compared overlapping sets of samples from two periods for *Ptros* and selected demographic parameters, including numbers of mussels aged 4-7 years (*Ptros* were calculated for mussels in this age group) using the Wilcoxon signed-rank test.

### Collection and analysis of historical data on Murman mussels

We searched for historical data on mussels from a 50-kilometer stretch of the coast with a center in the Tyuva Inlet (**Fig. 1b**). The choice of its boundaries was rather arbitrary, based on the assumed similarity of this coastal stretch to the Tyuva Inlet in terms of environmental conditions. We looked for any published data on Murman mussels comparable to our own data on the Tyuva, primarily for the data on mussel abundance estimated along vertical transects. We used Google Scholar with the keywords “*Mytilus*” and “Kola Bay” or “Barents Sea” to search for the recent literature and the catalogs of the Library of the Russian Academy of Sciences to search for older literature. We also used our own unpublished data on the abundance of mussels in the Klimkovka and the Dolgaya Inlets (**Fig. 1b**; **STable 2**). The mussels of each of these inlets were characterized using numerous samples collected at different depths along three littoral-sublittoral vertical transects in different parts of the inlet. A GAM model (normal distribution) with the Year as a predictor was used to analyze the long-term dynamics of mussel abundance based on all material. The dependent variable was transformed (log(x+1)). To compare the data on the abundance of mussels in different periods of time, the Mann-Whitney test for medians and the Wilcoxon Test for matched samples were used.

## Results

### Prediction of taxonomic composition by morphotypes

Proportion of *M. trossulus* genotypes (empirical *Ptros*) in genotyped samples from the Tyuva ranged from 0.06 to 0.90, i.e., from almost pure *ME* to almost pure *MT*. The proportion of T-morphotypes (*PT*) in the same samples ranged from 0.08 to 0.98. The relationship between the two indices was close to proportionality (**Fig. 2**). Parameters of regression models describing the dependence of empirical *Ptros* on *PT* and Dataset (i.e. sets of samples from the Tyuva, from brackish and saline habitats in the Barents Sea) are given in **STable 3**. The empirical data from the Tyuva generally agreed well with the model predictions, but some samples apparently had too low or too high *Ptros* at a given *PT*, which may be partly due to generally small sample sizes. The regression line corresponding to the Tyuva occupied an intermediate position between the lines corresponding to the other two sample sets but was closer to saline habitats (**Fig. 2** insert).

**Figure 2.**
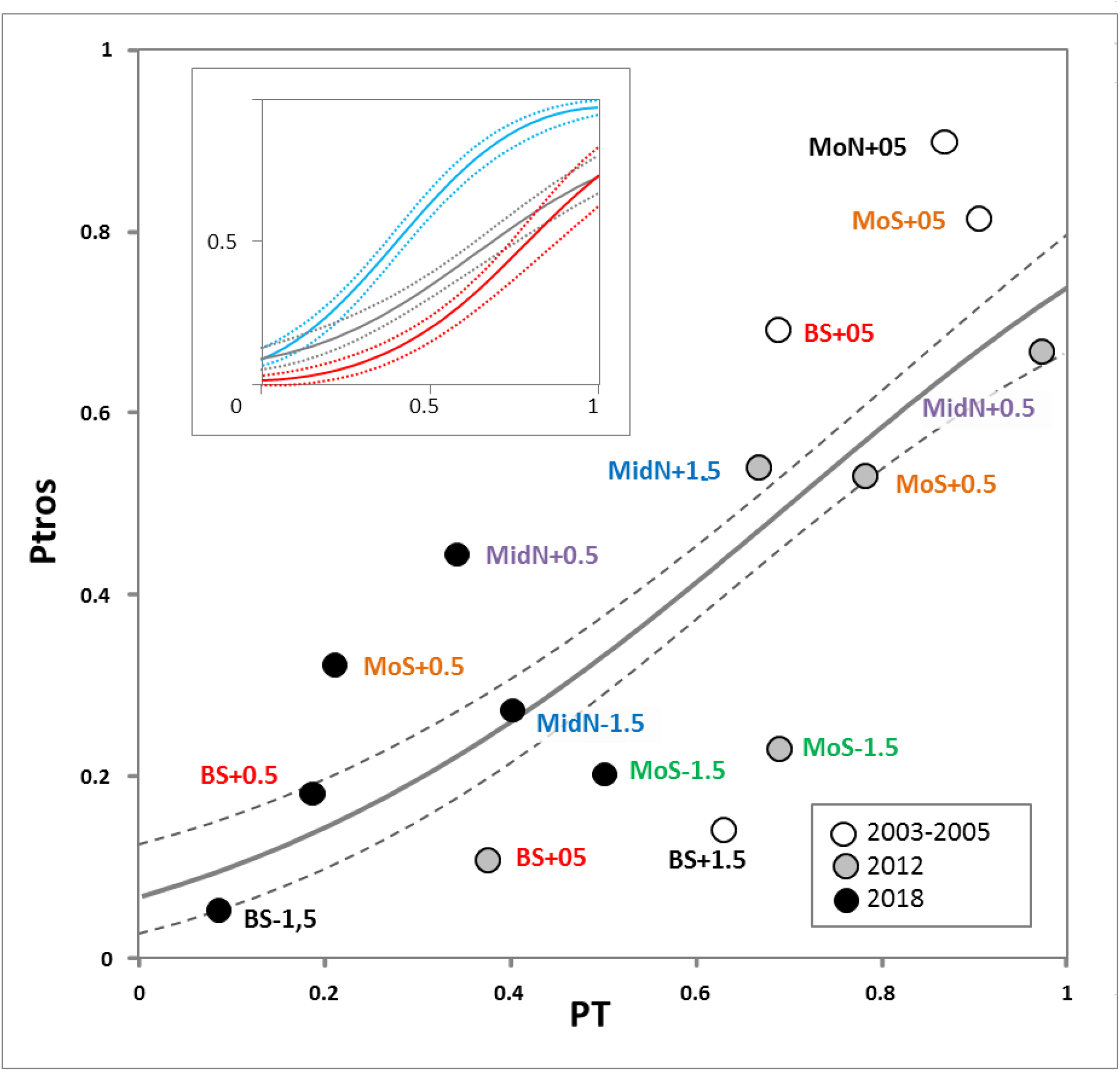
Dependence of proportion of *M. trossulus* genotypes (*Ptros*) on proportion of T-morphotypes (*PT*) in samples from the Tyuva. Dots are empirical estimates, color reflects the time period of sampling (see the legend). Sampling localities are labeled as in **Fig.1f**, repeated samples from the same localities are highlighted in font color. Solid line is regression model predictions, dashed lines are boundaries of 95% confidence interval of regression. The same regression is shown in the insert together with the corresponding regressions for samples from brackish (<30 ppt; blue) and saline (>30 ppt; Tyuva samples not considered; red line) localities in the Barents Sea from the study of Khaitov et al., 2021.

Since the genotyped collections from the Tyuva included samples taken from the same settlements at various time points, we give the first idea of the scale and direction of *Ptros* temporal dynamics in **Fig. 2**. In general, the proportion of *MT* decreased with time. For example, *Ptros* in BS+05 (the littoral part of the Bed) was 0.69 in 2004 and 0.11 in 2018, while *Ptros* in MoS+05 (rocky littoral) 0.81 in 2005 and 0.33 in 2018. The differences between the collections made at the same time points from different depths are also noteworthy: *Ptros* was always higher in the littoral than in the sublittoral (by 12-30%, on average by 16%).

### Tyuva mussels and their environment in 2009-2010

Salinity at sampling sites varied broadly during the tidal cycle, especially in the upper part of the inlet (4-34 ppt, with minimal values at low tide). According to the fitted model (**STable 4**), the predicted salinity increased with the distance from the river and was on average 16 ppt at the top of the inlet and 23 ppt at its entrance (**Fig. 1c**). The amplitude of predicted salinity fluctuations during the tidal cycle was maximal at the top (1-29 ppt at R+05) and minimal at the entrance (18-27 ppt at MoN).

The other environmental parameters of the sampling localities are provided in **STable 1**. Their variation generally corresponded to the literature data and visual observations (see Materials and Methods): the transects at the top of the inlet were wide, because the shore was gently sloping, especially in the south, and sandy. The transects at the mouth and in the middle of the inlet were narrow, because the shore was steep, rocky, and abundantly overgrown with fucoids on the littoral and kelps in the sublittoral (**Fig. 3**).

**Figure 3.**
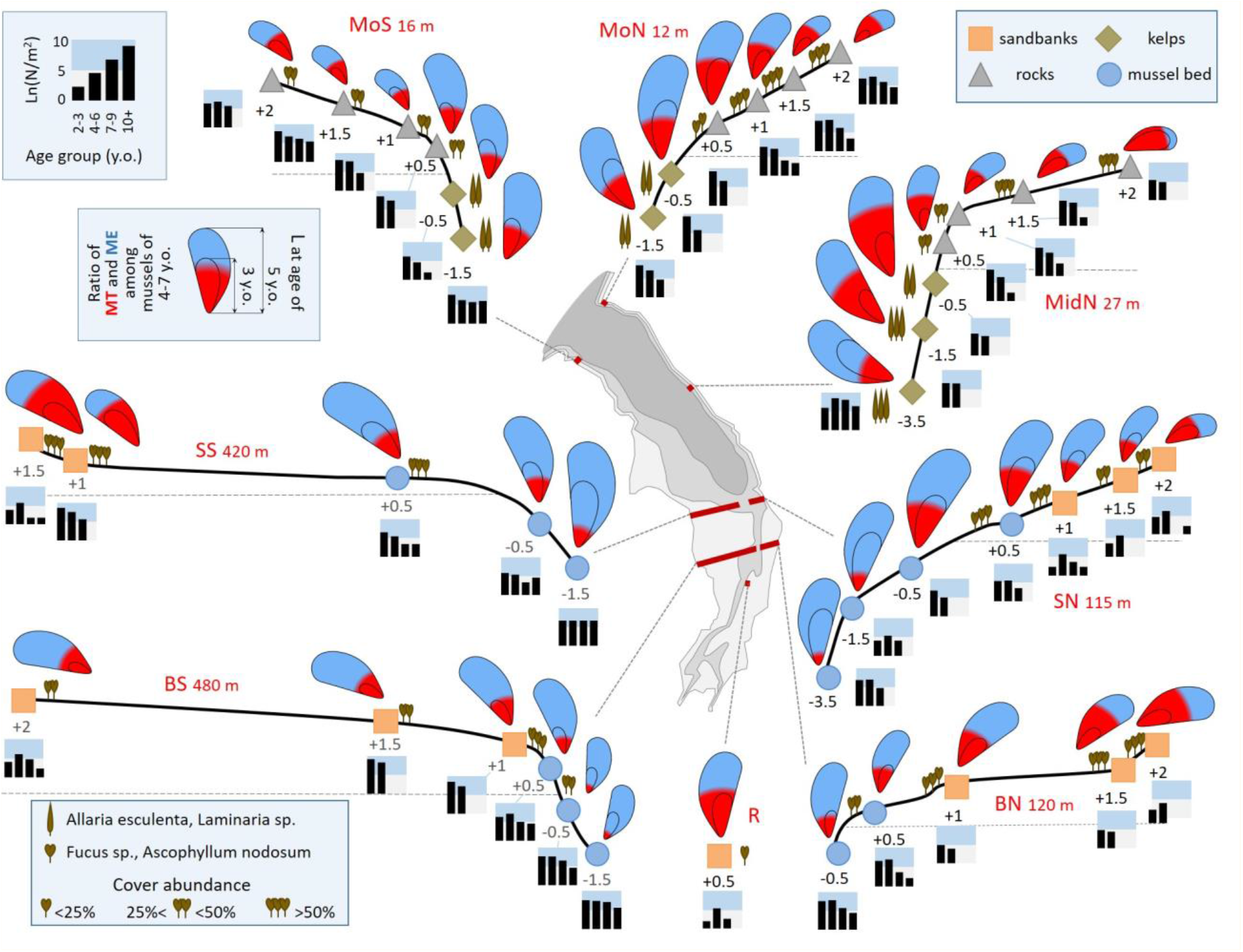
Characteristics of mussel settlements in the Tyuva Inlet in 2009-2010. Schematic profiles of transects where mussels were studied are given (note that the width of different transects is given on a different scale because the steepness of the shore differed greatly). The transect width is specified. Dots denote studied settlements, the depths in meters from the chart datum are indicated. The color of the dots reflects the habitat (see the legend). The icons showing algae represent the dominant algal species and their cover abundance, as indicated in the legend. The size of mussels in the pictograms is proportional to the average size of five-year-old mussels, the “annual ring” is proportional to the average size at the age of three years, while the color filling is proportional to the ratio of *MT* (red) and *ME* (blue) among mussels aged 4-7 years. Histograms show age structures, the logarithms of the average density of mussels of different age groups per m^-2^ are given (see the legend). Other notations are as in **Fig. 1c**.

Demographic parameters varied broadly between settlements, e.g. *N* from a few tens to tens of thousands ind*m^-2^, *W* from tens of grams to as much as ten kilograms m^-2^, *L5* from 17 to 38 mm. The largest mussel found was 87 mm in length. Predicted *Ptros* varied in a range of 0.10-0.73 (**STable 1**). The patterns of spatial variation of *Ptros* and some demographic characteristics can be deduced from **Fig. 3**. In terms of the total abundance, the greatest differences were registered between very sparse settlements on the sandbanks and dense settlements in the rest of the inlet. In terms of the age structure, the differences between the transects through rocky littoral and kelp forests (MoN, MoS, and MidN), where juveniles were dominating, and the transects in the upper part of the inlet, where there were few juveniles, are noteworthy. The average size of mussels of the same age increased consistently with the depth along all transects, except those through the densest part of the Bed (BS, BN), where an opposite trend was observed. Predicted *Ptros* generally decreased with the depth, but there were two striking deviations from this general pattern. Firstly, an anomalously high *Ptros* was recorded at MoN-0.5 and MidN-0.5. Secondly, *Ptros* was low throughout the Bed. It was noticeably lower there than on the sandbanks and at the same-depth horizons of the other transects (**Fig. 3**).

CCA was used for an analysis of associations between all environmental, demographic and taxonomic parameters (**Fig. 4**). Out of the eight initially considered environmental parameters, only three were included in the optimal CCA model: *Depth, Distance* and *Slope*. The influence of *Distance* and *Depth* on the ordination was significant while that of *Slope* was not (**STable 5**). Two first canonical axes were statistically significant, explaining 41.1% of the total inertia. CCA1 showed a high positive correlation with *Distance. Depth* and *Slope* were more associated with CCA2; positive values of CCA2 correspond to the littoral zone. Among the demographic parameters associated with CCA1, *N* and *N2-3* showed a positive association, whereas *B, N4-6, N7-9, N10, GI, L5, Lmax* showed a negative association (**Fig. 4**). This means that mussels, especially young ones, were more numerous in the settlements of the outer part of the inlet compared to the settlements of the inner part of the inlet, but also that mussels in the former settlements were slow-growing and their total biomass was not large. *Ptros* demonstrated the highest positive correlation with CCA2: that is, the proportion of MT on the littoral was higher than in the sublittoral. Notably, the parameters of size (*GI, L5*, and *Lmax*) also tended to be positively related to CCA2.

**Figure 4.**
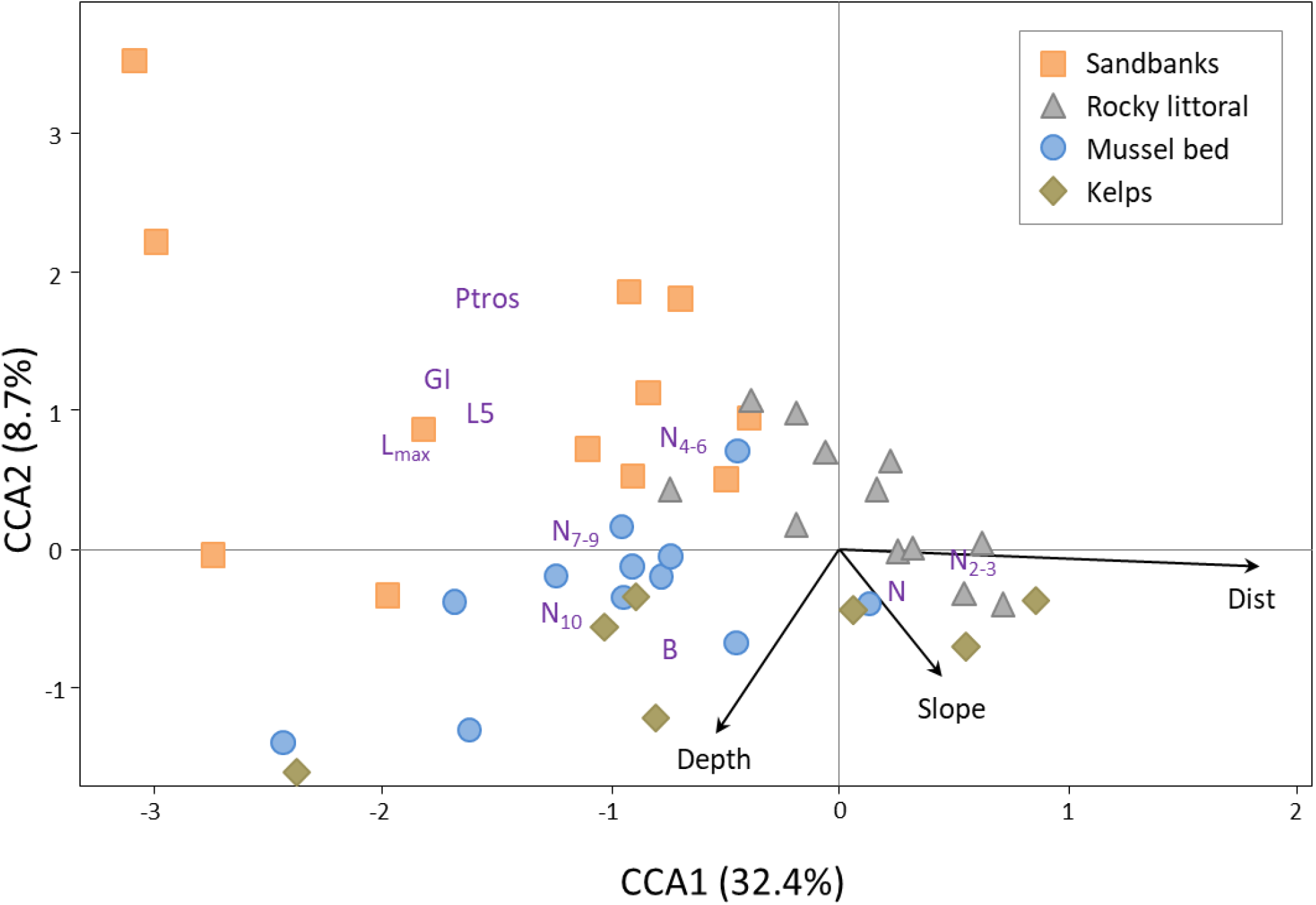
Canonical symmetrical correspondence analysis (CCA) ordination of mussel settlements by demographic and taxonomic parameters in 2009-2010. Each point represents a mussel settlement; settlements from the kelp forests, the rocky littoral, the sandbanks, and the mussel bed are shown with points of different form and color (see the legend). Text markers represent demographic and taxonomic parameters. Arrows indicate environmental constraints (direction and length of arrows show the degree of association between canonical axes and constraints).

Considering that these parameters were negatively related to CCA1, these observations may mean that mussels on the broad sandbanks in the top of the Tyuva Inlet were relatively larger (**Fig. 4**). Settlements from different habitats showed a tendency to nonrandom ordination in CCA. Settlements of sandbanks (upper left quadrant of triplot, high *Ptros*, low *N*, deficit of juveniles, large mussel sizes) were particularly strongly separated from the others.

PERMANOVA followed by pairwise comparisons of settlements from different habitats revealed that the settlements of sandbanks differed strongly from all the others and that the differences between the mussel bed and rocky the littoral were marginally significant after correction for multiple testing (uncorrected p=0.02, **STable 6**).

Analysis of associations between *Ptros* and environmental predictors using GLMM (**STable 7**) showed that only two variables significantly influenced the taxonomic structure of the settlements: *Depth* (the greater the depth, the lower the *Ptros*) and *Exp* (*Ptros* was higher on the north coast).

### Temporal changes in Tyuva settlements in 2004-2018

The first and the second axes of the CA based on the matrix of demographic parameters in all the material studied (**Fig. 5a**) explained 57.2% and 20.8% of the total inertia, respectively. The same set of parameters was associated in a similar manner with CA1 and CCA1 in a separate analysis of 2009-10 data, but the ordination of samples from different habitats along CA1 and CCA1 was different (compare **Fig. 4** and **Fig. 5a**)). The reason behind the differences is the large-scale temporal dynamics, which can be inferred from **Fig. 5a**, where samples from different time periods are highlighted, and from the results of the GAM regression analysis. It is striking that all the 2004-2005 samples (highlighted in black) in the figure are centered further to the left of the graph than most samples from other periods. The differences in the ordination of 2009-10, 2012, and 2018 samples are less prominent. According to GAM, over the entire observation period, there were significant changes in CA1 scores toward larger values (i.e., primarily a decrease in the number of adults and an increase in the number of juveniles) in the settlements from littoral rocks and from the Bed (**STable 8**; **SFig. 2**). The arrows in **Fig. 5a** show changes in BS+05 (littoral part of the Bed): the samples moved from left to right throughout the 13-year-long study period. These changes actually reflect a gradual “degradation” of the littoral part of the Bed as a dense settlement dominated by old mussels. Judging from visual observations, the change started already in 2010 but affected BS+05 somewhat later. By 2018, the littoral part of the Bed had completely disappeared(see **SFig. 3** for age frequency distributions in BS+05 and photos of the Bed at different years).

**Figure 5.**
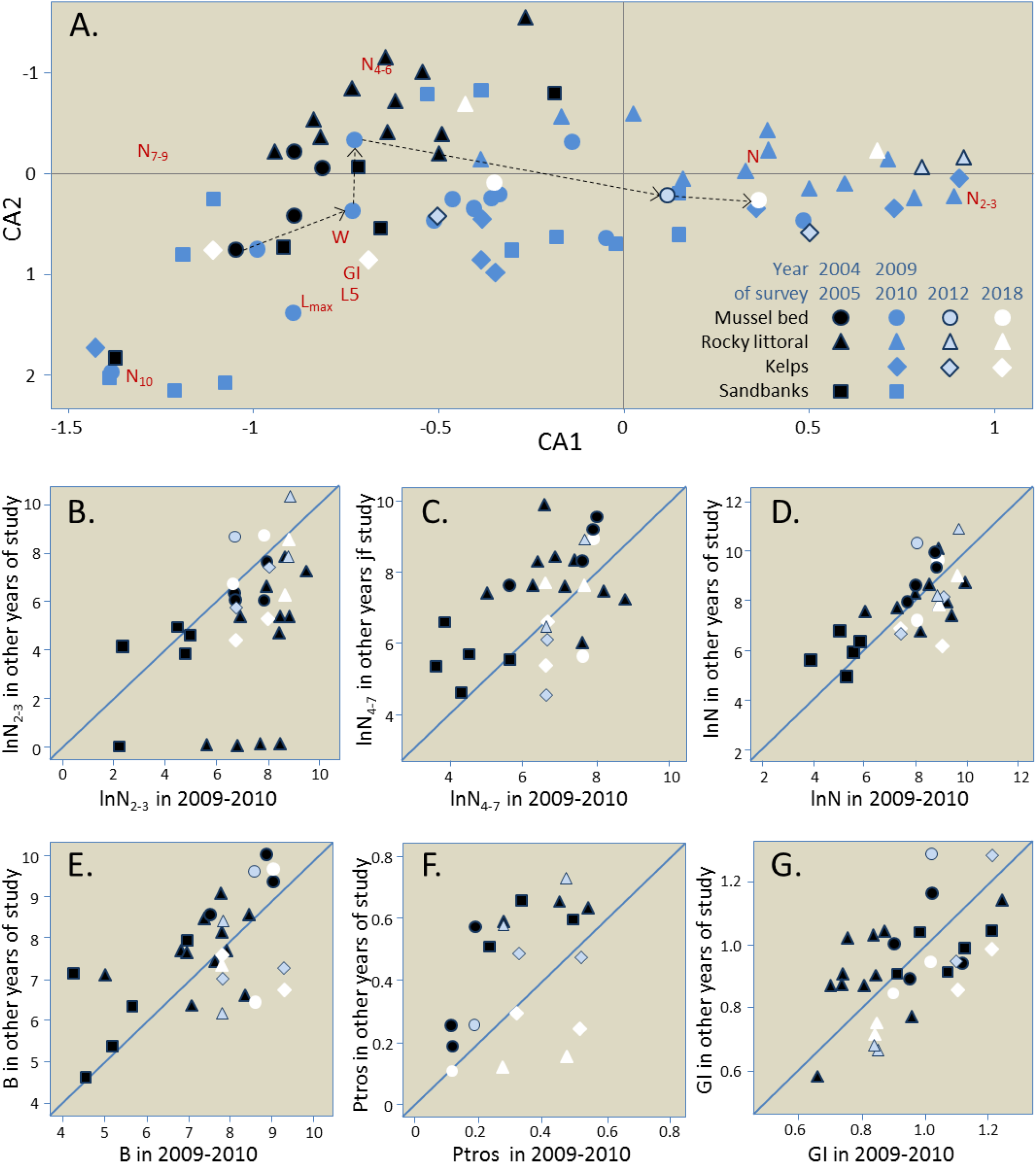
Temporal changes in demographics of the Tyuva mussels. (a) Correspondence Analysis (CA) ordination of mussel settlements by demographic parameters in all surveys (data on the same settlements in different years are considered independently). Demographic characteristics are indicated by abbreviated names. Settlements are marked with signs, settlements from different habitats are shown with signs of different shape, and those studied in different years are shown with signs of different color, as shown in the legend. Arrows show temporal changes in BS+05 investigated in 2005, 2009, 2010, 2012, and 2018. (b-g) Temporal changes in the repeatedly studied settlements by *N2-3* (b), density of mussels aged 4-7 years (c), *N* (d), *B* (e), *Ptros* (f), *GI* (g). All abundance values are logarithmic. All settlements were studied in 2009-10 (their characteristics in 2009-10 are plotted on the horizontal axis) and at one or more other time points (vertical axis). If the settlements did not change over time, the points lie on the diagonal.

Regression analysis of *Ptros* variation with time also revealed significant changes in the settlements from littoral rocks and in the Bed: in both habitats the proportion of MT decreased with time (**STable 9, SFig. 2**). In settlements from sandbanks and kelps, the tendency was the same, but insignificant (**STable 9, SFig. 2**). This could be due to the scarcity of data on these habitats.

**Fig. 5b-g** illustrates the temporal variations of *Ptros* and selected demographic parameters in settlements studied both in 2004-05 and in 2009-10. Changes are the most noticeable in *Ptros* (**Fig. 5f**) decreased by 22% on average; Wilcoxon test: uncorrected p=0.0039), *N2-3* (**Fig. 5b**, increased fivefold; p=0.00013), *N4-7* (**Fig. 5c**, decreased by a factor of three; p=0.011) and *B* (**Fig. 5e**, decreased twice; p=0.0042). Analysis of changes using PERMANOVA confirmed the unidirectional change in all habitats because no significant interaction between *Period* and *Habitat* factors was revealed (**STable 10**). SIMPER procedure showed that densities of mussels of different age groups made the greatest contribution to the change (in total, 72% of the cumulative contribution, **STable 11**).

### Long-term dynamics of the Murman mussels

Most of the old studies provide biomass data (**STable 2**), so these were the only data that we analyzed. In total, we found 34 estimates of average mussel biomass on vertical transects throughout the littoral, obtained in 1933-2002 and representing 18 coastal sites. Most of the sites represented individual inlets, including the Tyuva, and, with one exception (Ura Inlet, 1961), the site was characterized by one transect at a time (**STable 2, Fig. 6a** and caption to the figure). Therefore, to analyze the temporal dynamics, we averaged the data for this inlet and used the average value of biomass per site per study. These data together with comparable data from our studies are visualized in **Fig. 6b**, which also shows the long-term change in temperature in the Barents Sea. As can be seen from the **STable 2** and **Fig.6**, the two overlapping data sets are the most informative. (1) The data accumulated in the course of the VNIRO (Research Institute of Fisheries and Oceanography) monitoring surveys of commercial bivalves in 1960-1961 (16 sites), 1971 (7, all overlapping with 1961) and 1981 (3, all overlapping with 1961 and one also with 1971). (2) Data from five inlets that have been studied more than twice (including our own collections): Zelenetskaya Zapadnaya (studied in 1961, 1971, 1981 and 1985), Ura (1960, 1961, 1971, 2002), Tyuva (1960, 1961, 1971, 2004, 2009), Klimkovka and Dolgaya (both were investigated in 1961, 1971 and 2009). **Fig. 6b** shows that the values of the average biomass in 1960-1961, at the end of the 30-year-long period of high SST in the Barents Sea, were unprecedentedly high everywhere (range 2.0 - 32.0 kg*m^-2^, median 6.5 kg*m^-2^). When compared to 1960-1961, the values of the biomass in 1971 were almost an order of magnitude lower (range 0.1 - 2.1, median 0.7 kg*m^-2^). Differences between the two time sample sets were significant (Mann-Whitney test for medians, p=0.0002), and so were the differences between the 7 paired samples (Wilcoxon test for matched samples, p=0.02). In 2009, the year that was preceded by a 5-year-long period of anomalously high SST, the values of the mean biomass in the Tyuva, the Dolgaya, and the Klimkovka were still many times lower than in 1960-1961. Given all the data, the values of the biomass in the 2000s were on average slightly larger than in the 1970s and 1980s; the fitted GAM demonstrated the minimal biomass in 1980-1990 (**Fig. 6b**; **STable 12**).

**Figure 6.**
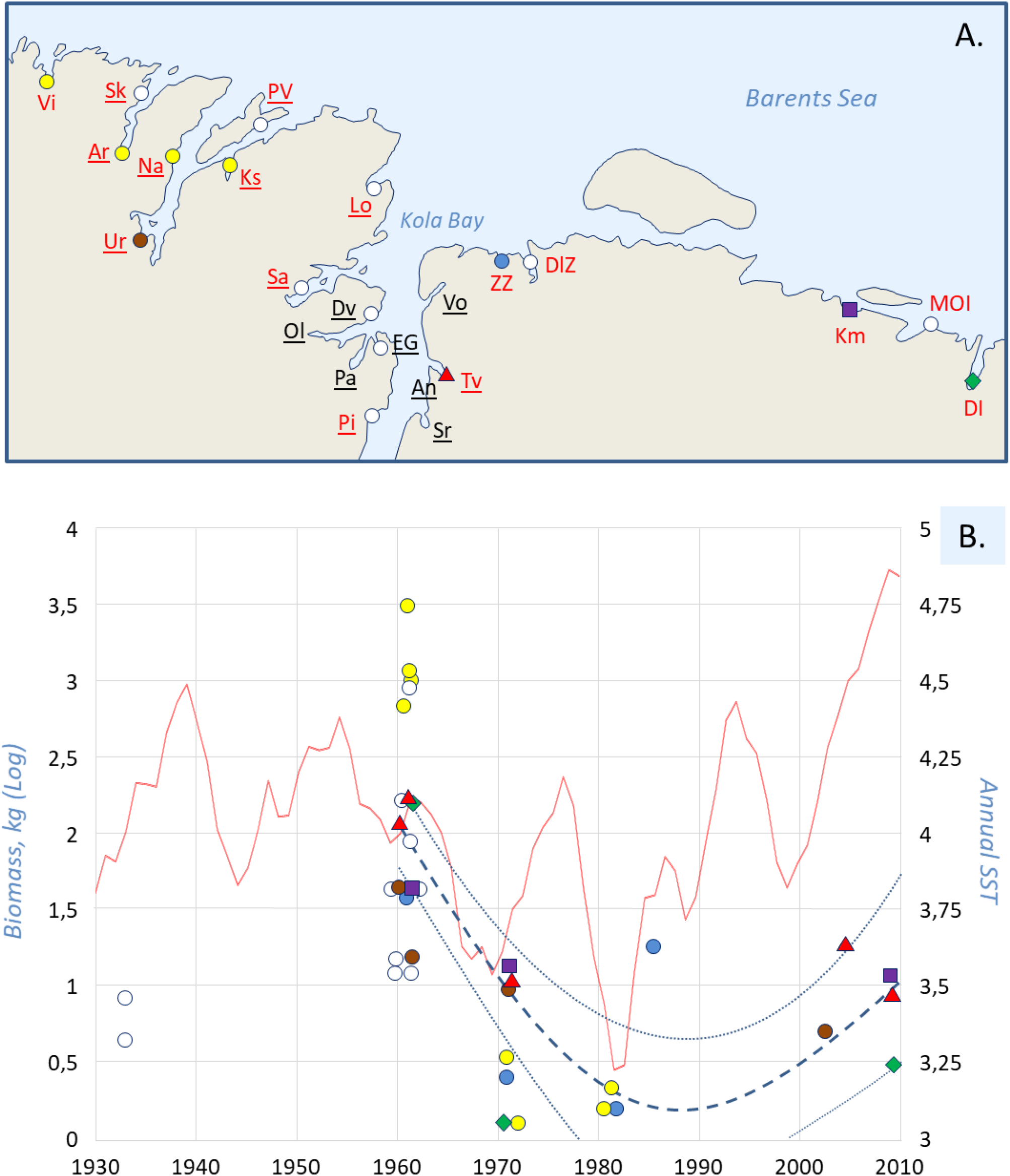
Long-term dynamics of mussels in the northern Kola Bay and its vicinity according to published data and our own data. (**a**) Map of historical mussel survey sites labeled as in **STable 2**. The following sites are mentioned in the text: Tv – Tyuva Inlet, Km – Klimkovka Inlet, ZZ – Zelenetskaya Zapadnaya Inlet, DL – Dolgaya Inlet, Ur – Ura Inlet. Sites where the mussel biomass was estimated on vertical transects through the mussel-inhabited zone of the littoral are marked with filled signs: white – the site was examined once in the history of studies, yellow – twice, other colors – more than twice (see **STable 2** for details). The names of the VNIRO monitoring study sites in 1960-1985 (Romanova, 1969; Antipova et al., 1984) are in red, and those of the sites from qualitative studies in the 1920s (Guryanova et al., 1928, 1929, 1930) are underlined. Other data are from Zatsepin et al., 1948; Kostylev, 1989; Milyutin & Sokolov, 2006 and our studies. (b) Temporal dynamics of water temperature and mussel biomass. The biomass is displayed on the left axis, while the temperature, on the right axis. Signs represent estimates of average biomass on vertical transects through the mussel-inhabited littoral zone at different locations, with data from different sites labeled as in map (a). Bold dashed line is regression model predictions for 1960-2009, thin dashed lines are boundaries of 95% confidence interval of regression. The red broken line is a five-year running mean water temperature in the Kola hydrological section (0-200 m; stations 3-7) (Bochkov 1982; 2005; www.pinro.ru).

Our data on the Tyuva, the Klimkovka, and the Dolgaya (**STable** 2) and the data on the Ura (Milyutin & Sokolov, 2006) indicate that in the 2000s the mussel abundance in the upper sublittoral down to a depth of 5 m was similar to that in the littoral. Unfortunately, there is nothing to compare these data with. Strange as it may appear, there are no older data on mussel abundance in the sublittoral even though in the monitoring studies of commercial bivalves by VNIRO in 1960s-1980s the upper sublittoral was examined for bivalves using the diving method. There are, for instance, data on abundance of sublittoral *Modiolus* for the Tyuva, the Klimkovka, the Dolgaya, and the Ura. However, sublittoral *Mytilus* are not even mentioned. It is noteworthy that most of the sublittoral collections in the Klimkovka, the Dolgaya, and the Tyuva were from kelp forests. Another remarkable circumstance is that the VNIRO study from the 1960s makes no reference to a mussel bed in the Tyuva, though mussel beds in some other sites are mentioned (Romanova, 1969).

Purely qualitative studies made in the 1920s by Guryanova, Zaks and Ushakov (1928, 1929) also provide some valuable information in the context of our research. These authors described the littoral at different parts of the coast (**Fig. 6a**), noting, in particular, characteristic mussel habitats (essentially the same as we identified in the Tyuva, see **SFig. 1** for an illustration of the habitats and **STable 2**). They also described the littoral communities at the top of the Tyuva Inlet in 1923 and provided a map (see **SFig. 3**). Though these authors did not observe the mussel bed where we found it in the 2000s, they did notice two relatively small mussel patches in that area.

## Discussion

We conducted a phenomenological study of the Murman mussels, using the Tyuva populations as an example, for the first time since the 1920s-1930s. The difference of our study from previous local ones (Guryanova et al., 1928, 1929; Zatsepin et al., 1948; Matveeva, 1948) is that we analyzed the interannual dynamics of the mussel settlements and took into account their taxonomic structure. An “old-fashioned” descriptive character and an emphasis on the taxonomic heterogeneity of the object distinguishes our research from the majority of modern ecological mussel studies, which are hypotheses-driven and often ignore the species identity of mussels (for a discussion of the latter issue, see Katolikova et al., 2016). By using a parsimonious morphological method of determining the taxonomic structure of the samples, we managed to map the species distributions on scales from tens of meters to several kilometers in unprecedented detail. We also extracted data on mussel abundance on Murman from the Soviet “gray” literature, inaccessible to a broad readership, and for the first time summarized the data on the long-term dynamics of subarctic mussels. Below we discuss first the distributional patterns of mussels, then their interannual and decadal dynamics, and, finally, the issue of *ME* and *MT*.

### Distributional patterns

The spatial patterns in mussel demography observed in the Tyuva during our large-scale surveys in 2009-2010 can be explained, as a first approximation, by the influence of abiotic factors, food availability, and density-dependent effects in the settlements. The following patterns can be outlined. 1) An almost ubiquitous distribution of mussels in the depth range from 4 m to +2 m at average surface salinity above 15 ppt; at lower salinity the mussels disappeared from the littoral. 2) The presence of the Bed in the river mouth in the top of the inlet, where the input of nutrients from the river and a fast water flow provide the best conditions for mussel feeding. 3) A trend toward increasing mussel size from the more wind-exposed mouth of the inlet, where it can be limited by wave action, to its top, which is more sheltered and where nutritional conditions are better. 4) An almost ubiquitous deficit of juveniles in the upper part of the inlet due to worse conditions for larvae and juveniles associated with strong salinity fluctuations and lack of substrate for settling on sandbanks, as well as on the Bed, where the space is occupied by large mussels. 5) A decrease in mussel size with littoral elevation associated with the negative impact of aerial exposure on their growth. At the same time, the relationship between the mussel size and the depth was inverted on transects across the riverine part of the Bed (BS, BN), where the mussels in scarce settlements on the sandbanks were on average larger than those in the Bed. This inverted relationship may be explained by the negative effect of the high density of mussels in the bed on their growth. This effect is more pronounced in the center of the bed and disappears towards its periphery (cf. Okamura, 1986).

Though all the patterns mentioned above are trivial, they are not always easy to identify in boreal seas because of the confounding pressure of mussel predators (Seed & Suchanek, 1992). This pressure seems to be weakened in the Tyuva Inlet, where the only major enemy of mussels are common eiders (Krasnov & Goryaev, 2013). There are no littoral crabs in the Murman waters (Zatsepin et al., 1948). Starfish, dog whelks and sand shrimps occur in the region but we did not encounter any of them in the Tyuva. Finally, the mussels in the Tyuva are not harvested by humans.

The patterns of mussel habitat distribution identified in the littoral of the Tyuva Inlet in our study generally match those recorded in the 1920-1930s (Guryanova et al., 1928, 1929; Zatsepin et al., 1948; **SFig. 1**). It is as if time stood still for mussels in the Barents Sea littoral for a hundred years. However, this was definitely not so in the sublittoral. We found fairly numerous populations of fast-growing mussels in kelp forests. No such populations have been noted in the 20th century. In particular, they have not been recorded in the sublittoral studies of commercial bivalves of VNIRO in the 1960-1980s (Romanova, 1969; Antipova et al., 1984) either in the Tyuva or in any other inlets where we observed them in our study. No *Mytilus* populations are mentioned in studies of the kelp communities in the areas of the Kola Bay adjacent to the Tyuva in the beginning of the 20th century (Derjugin, 1915; Guryanova, 1924). Yet in a recent study of kelp communities from the same area the mussels were mentioned as an important component (Pavlova et al., 2018). One gets the impression that mussels have inhabited the kelp forests only in the recent decades.

Populations in kelp forests seem to be characteristic of Arctic mussels, for which the littoral poorly accessible due to small celestial tides, abrasive action of ice and extreme winter temperatures (Feder et al., 2003; Leopold et al., 2019; Sukhotin et al., 2008). They were also described in more temperate seas (e.g. British Isles, Connor, 1997; Gulf of St. Lawrence, Bégin et al., 2004; Aleutian Isles, Stewart & Konar, 2012). It is debatable whether kelp forests are suboptimal habitats for mussels, which they colonize when other habitats are scarce. It has been experimentally proven that, if there is an alternative, the larvae of the White Sea mussels avoid settling on or near kelps, probably due to the repellents released by the algae (Dobretsov, 1999; Dobretsov & Wahl, 2001). Indeed, mussels are rarely found in kelps in the White Sea (Plotkin et al., 2005). On the other hand, in the Gulf of St. Lawrence, the kelp canopy has been shown to promote successful mussel recruitment (Bégin et al., 2004). However that may be, the seeming absence of the Murman mussels in the kelp forests in the 1920s-1980s and their appearance there in the 21st century is intriguing. We will return to this mystery in the section on ME and MT (see below).

### Interannual dynamics

The most salient features of the temporal dynamics of the Tyuva mussels recorded in our study were synchronous changes in the age structure of settlements across the Tyuva between 2004-05 and 2009-10 and the increasing “degradation” of the littoral part of the Bed in 2010-2018. Between 2004-05 and 2009-10, the settlements became significantly “younger” everywhere. There were very few young (1-3 year-old mussels) in 2004-05, indicating poor recruitment (or poor survivorship of young mussels) in the early 2000s. In 2009-10, there were few old individuals born in the early 2000s, but many young individuals.

The fact that the changes were pervasive suggests a common causal factor in the dynamics, but we cannot say with certainty what factor it was. We do know that the annual SST has been increasing since the late 1990s (**Fig. 6b**), which means that the mass recruitment occurred in warmer years. However, this might have been a coincidence. According to the data from other mussel studies, synchronicity in the interannual dynamics of their settlements on a spatial scale comparable to the Tyuva Inlet is an exception rather than the rule (Stillman et al., 2000; Folmer et al., 2014; Khaitov & Lentsman, 2016; Khalaman et al., 2020; but see Westerbom et al., 2021 for an opposite example: a high year-to-year variation in recruitment of mussels related to salinity fluctuations in the mesohaline environment). The classic hypothesis suggested based on the data on the Wadden Sea mussels and other littoral bivalves explains the recruitment synchrony in their populations by the fact that the abundance of invertebrate predators feeding on spat is reduced during severe winters (Beukema et al., 2015 and references therein). Taking into account a very different thermal regime and the lack of such predators in the Tyuva we doubt that this hypothesis can explain our results.

Our choice of the Tyuva Inlet as a study site was partly due to the presence there of a mussel bed with an area of several hectares (Bufalova et al., 2005; this study). According to anecdotal evidence from local residents, the Bed had existed, seemingly unchanged, for at least 10 years before the start of our research. In 2010, visual observations indicated an incipient degradation of the littoral part of the Bed. What had been a solid carpet of mussels, mostly large ones, was gradually turning into a graveyard of their dead shells. By 2018, the degradation of the littoral part of the Bed has been completed, though there were plenty of juveniles at monitoring point BS+05 in 2012 and it had looked like the Bed should revive. In occasional studies in the Tyuva in the 20th century (Guryanova et al., 1928; Romanova, 1969; Antipova et al., 1984), no one noticed any large mussel bed there, which indicates the unstable nature of the Bed described in our study.

Mussel beds are known to exhibit large-scale dynamics, similar to what we observed in the Tyuva, which may be due to both “endogenous” and “exogenous” factors (Dankers et al., 2001; van der Meer et al., 2019; Khaitov & Lentsman, 2016 and references therein). Endogenous factors are associated with density-dependent processes in the bed itself: adult mussels prevent the recruitment of juveniles, and mass recruitment occurs only after the death of most of the old individuals. External factors are associated with physical disturbance such as storms, ice scouring, and cold waves in ice-free winters. Our observations were sketchy, and we do not know which of the factors were at play in the Tyuva.

### Long-term dynamics

Considering historical data on the Murman mussels, we may be fairly sure that their littoral populations collapsed between 1960 and 1970, having decreased in terms of biomass by an order of magnitude. These conclusions are mostly based on the VNIRO data (Romanova, 1969; Antipova, 1984, **Fig. 6**), which were probably obtained by comparable methods and therefore seem plausible. Estimates of littoral mussel abundance in the 1960s (median biomass 6.5 kg*m^-2^, **Fig. 6**) seem excessively high as compared with the rest of the data on the Murman mussels used in our paper. Nevertheless, they are not completely unrealistic. Matveeva (1948) reports a similar mussel biomass in 1939 for the eastern Murman site, located outside our study area and therefore not included in our search. We also know that in the subarctic Sea of Okhotsk the abundance of mussels is at present comparable with that in Murman in the 1960s (Ivanova & Tsurpalo, 2011; Khalaman et al., 2020). The available data suggest that by the beginning of the 21st century the populations had not fully recovered from the collapse of the 1960s (**Fig. 6**).

In recent decades, mussel populations (mainly *ME*) in boreal seas have shown a downward trend in abundance. The main hypotheses attempting to explain this phenomenon are associated with overfishing and the effect of warming climate, direct or indirect (e.g. through increased predator pressure) (Sorte et al., 2017; Baden et al., 2021 and references therein). Climate warming also explains the shift of the southern limit of the *ME* range northward in the western Atlantic (Jones et al., 2010). However, in the subarctic Barents Sea, the opposite relationship between the temperature and the mussel abundance and distribution is expected. Paleontological evidence suggests that during the warm periods of the Pleistocene-Holocene, mussel abundance in the Barents Sea region increased and their distribution area expanded deep into the Arctic (Hansen et al., 2011; Mangerud & Svendsen, 2018). The best example is the reappearance of mussels in Spitsbergen in the early 2000s, after an absence of a thousand years (Berge et al., 2005). The available data on mussel dynamics in Murman in the second half of the 20th century also agree with the hypothesis of a direct relationship between mussel abundance and water temperature. High biomasses were observed in 1960-61 at the end of a roughly 40-year-long period of predominantly high temperatures, and the subsequent collapse coincided with the beginning of a severe cold snap that lasted into the late 1990s, when a very warm spell, which we are still observing now, has started (Drinkwater, 2011; Fig. 6b). It is assumed that the whole ecosystem of the Barents Sea changes with the climate (Matishov et al., 2012; Ingvaldsen et al., 2021), although it is difficult to disentangle the effects of climate change from those of fishing for its most studied components such as sublittoral benthos, zooplankton and commercial fish species (Denisenko, 2001; Johannesen et al., 2012). In accordance with the temperature and the correlated temporal variation of primary production, the general trend in biomass for boreal species in the Barents Sea in 1950–2013 was U-shaped, with low values in the 1960–1980s (Pedersen et al., 2021). Against this background, a weak response of the Barents sea littoral mussels to the warming in the early 21st century seems unusual. This is another mystery of the long-term dynamics of the Murman mussels, and we will return to it below.

### *Mytilus edulis* and *M. trossulus*

We could not directly identify the contribution of *ME* and *MT* to the demographic structure of settlements because the morphological method used in our study did not allow the species assignment of individuals and was applicable only to mussels aged 4-7 years. From an earlier study by Bufalova et al. (2005), however, we know that there are no differences in the growth rates of *ME* and *MT* in the Tyuva. Therefore, we can only discuss how these species divided space and how their relative frequencies in populations varied with time in 2004-2018.

In the Tyuva, *ME* and *MT* inhabited essentially the same habitats as they do in allopatry. In particular, mass populations in kelps have been described both for *ME*, e.g., in the Pechora Sea (Sukhotin et al., 2008), and for *MT*, e.g., in the Aleutian Islands (Stewart & Konar, 2012). At the same time, these two species partially shared space and habitats with each other in the Tyuva. Their distribution was fairly regular (the deeper the more *ME*; an excess of *ME* on the Bed), although elements of mosaic distribution could also be seen (**Fig. 3**).

In their early review on sympatric *ME* and *MT*, Riginos & Cunningham (2005) compared the two zones of their sympatry known at the time, at the entrance to the Baltic Sea and in the Canadian Maritimes (Western Atlantic), and pointed out striking differences in the habitat distribution of these species in the two zones. In the former, their distribution is governed by salinity, with *MT* thriving in the extremely freshened environments of the central Baltic. In the more oceanic habitats of the Western Atlantic, these two species are distributed mosaically, with patches dominated by different species alternating at a scale of kilometers – tens of kilometers; *MT* tends to dominate in more exposed sites and *ME*, in more sheltered ones. If there is a relationship between the distribution of these two species and salinity (and the degree of wave exposure, which is difficult to separate from salinity) in Western Atlantic, it is the opposite to that observed in the Baltic. Riginos & Cunningham (2005) raised the question of whether the differentiation of ecological traits between species in the sympatry reflects their ancient divergence in the allopatry or whether it evolved already in the sympatry as a result of competition. This question seems to be unresolved to this day. Nevertheless, one might expect similarities in the habitat distribution of the species in the Barents Sea and in the Western Atlantic, given the similar salinity regimes and a probably very recent origin of the Barents Sea *MT* from the Western Atlantic (Väinölä & Strelkov, 2011, see below). Indeed, there is clearly no positive correlation between salinity and *MT* proportion in the settlements either in the Western Atlantic or in the Tyuva Inlet, but there is a tendency (not significant in the Tyuva) for *ME* to be more frequent in sheltered localities and for *MT*, in the exposed ones (Bates & Innes, 1995; Comesaña et al., 1999; Tam & Scrosati, 2014).

As for the segregation of these species by depth in the Western Atlantic, no one has studied it in detail on vertical transects as we have, which makes direct comparison difficult. No consistent differences were shown between settlements from the lower and the middle intertidal levels in the Canadian Maritimes (Moreau et al., 2005). Based on the re-analysis of published data, Riginos & Cunningham (2005) suggested that *ME* could be more common in the sublittoral than in the littoral. Further, it has been shown that *ME* larvae settle on average deeper than *MT* larvae, both in the laboratory and in the field (Kenchington et al., 2002; Freeman et al., 2002), which may result in an uneven distribution of species by depth. Consistent with this observation is the fact that *MT* mussels are more likely to occur at shallower depths on ropes of suspended mussel aquaculture than *ME* in the contact zone in Scotland (Michalek et al., 2021 and references therein). To note, segregation between competing mussel species by depth has been repeatedly observed in pairs other than *ME* and *MT* such as *MT* and *M. galloprovincialis* in California, Schneider & Helmuth, 2007) and *Perna perna* and *M. galloprovincialis* in South Africa (Bownes & McQuaid, 2006). In these cases, the competitors were a native species and a recent invasive species that partially displaced the native species from its intertidal habitat (a situation probably similar to *ME* and *MT* in the Barents Sea, see below).

There are a few distinctive features of *ME* and *MT*, which were left out in our study but may explain their distribution in the Tyuva. In the White Sea littoral, *MT* is more often found on algal substrates, while *ME* is found on bottom substrates (Katolikova et al., 2016). If segregation by substrate is the same in the Tyuva, this may explain the increased numbers of *ME* on the Bed, where algal substrates are scarce, compared to other littoral sites from the same depths. *ME* and *MT* differ in their aggregation behavior, with *ME* generally aggregating better (Liu et al., 2011). This behavioral feature can also be an advantage for *ME* in the Bed, where mussels form large aggregations.

The ratio of *MT* and *ME* in the Tyuva Inlet changed significantly not only in space, but also in time, the changes being synchronous across the inlet. There was a decreasing trend in the proportion of *MT* throughout the observation period. Between 2004-05 and 2009-10, *Ptros* decreased everywhere, by 22% on average. Again, the only factor which seems to be correlated with this change is the mean annual temperature, which increased during the study period. Indeed, *MT* is a more stenothermal species than *ME* (Rayssac et al., 2010; Fly & Hilbish, 2013). In field experiments in the White Sea, at water temperatures above 16°C adult *MT* have shown an increased heart rate and hence a poorer physiological performance than *ME* (Bakhmet et al., 2022). The negative effect of rising temperatures on *MT* has been considered as a possible factor explaining the replacement of *MT* by *ME* in the Oresund Strait at the entrance to the Baltic Sea between 1987 and 2005 (Strelkov et al., 2017). It is obvious that in an inlet of the Barents Sea in hot weather the littoral at low tide as well as the shallow waters can occasionally warm above 16°C. On the other hand, it is unlikely that temperature is as critical for *MT* in the Barents Sea (latitude 69’) as it is in the Oresund (56’), which lies at the southern boundary of its distribution in continental Europe.

Unfortunately, we cannot travel back in time to the 20th century, when the mussels demonstrated large-scale dynamics, and find out how it was related to their taxonomy. We can only speculate on this issue. Väinölä & Strelkov (2011) once hypothesized that *MT* invaded the Barents Sea during World War II with Allied convoys from the western Atlantic and established stable populations there after the 1960s, when a window of opportunity opened for them after the collapse of the native (*ME*) populations. Their hypothesis was based on, firstly, an increased incidence of *MT* in port areas and, secondly, genetic similarities between *MT* populations in the two regions. A recent genomic study has confirmed the similarity between *MT* populations in the Kola Bay and the Gulf of St. Lawrence as well as a young age of the hybrid zone between *ME* and *MT* in the Kola Bay, where the gene pools of hybridizing species do not bear any traces of recent introgression (Simon et al., 2021).

If the hypothesis of Väinölä & Strelkov (2011) is true, we can assume that in the second half of the 20th century the “common mussel” system went from a single-species (*ME*) state to a two-species state (*ME* and *MT*), and its parameters changed. The two mysteries of the century-long mussel dynamics in Murman described above, the expansion of mussels into kelp forests between 1970s and 2000s and a weak response of the mussel populations to the current climate warming, can theoretically be explained by this systemic transition. Perhaps the invasive species, better adapted to the littoral conditions, competitively displaced *ME* from the littoral to the sublittoral, where it colonized a previously unoccupied suboptimal habitat, kelp forests. A system of two competing and hybridizing species with different temperature preferences is unlikely to respond to climate changes in the same way as single-species populations. On the one hand, such a system may be less susceptible to climate fluctuations, as one species gains an advantage during cold periods and the other during warm ones. On the other hand, competition and hybridization should negatively affect the fitness of both species, reducing the growth potential of their populations. The latter consideration might explain a weak response of the Murman mussels to the current warming.

## Data availability

Individual data on genotype, age, and morphotype of genotyped mussels from the Tyuva Inlet are deposited in the database of St. Petersburg State University (http://hdl.handle.net/11701/38590). Information on quantitative samples of the Tyuva mussels is deposited ibid.

## Author Contributions

JM, PS, MS, MK, SM and VK contributed to conception and design of the study. PS, MK, MS, SM, LB, JM and MG organized mussel sampling. PS, JM, MS and MK provided mussel genotyping. JM and VM identified mussel morptotypes. JM and MS determined age and size of mussels. JM and EG conducted a search and analyses of historical literature. JM curated the data. VK, JM and PS performed the statistical analyses. PS, JM, VK and MK drafted the manuscript. All authors read, approved, and contributed to the final manuscript.

## Acknowledgments

We would like to thank all participants of our expeditions to Tyuva, especially Sergey Goldin, Sergey Shtinnikov, Dmitry Redkin, Helena Bufalova, Gita Paskerova, and Helena Shoshina for their help in the fieldwork, St. Petersburg University Research Park (https://researchpark.spbu.ru/) for assistance in laboratory analyses and Natalia Lentsman for English language editing of the manuscript.

