## Supplementary figures for "Taxonomically mixed blue mussel *Mytilus* populations are spatially heterogeneous and temporally unstable in the subarctic Barents Sea"

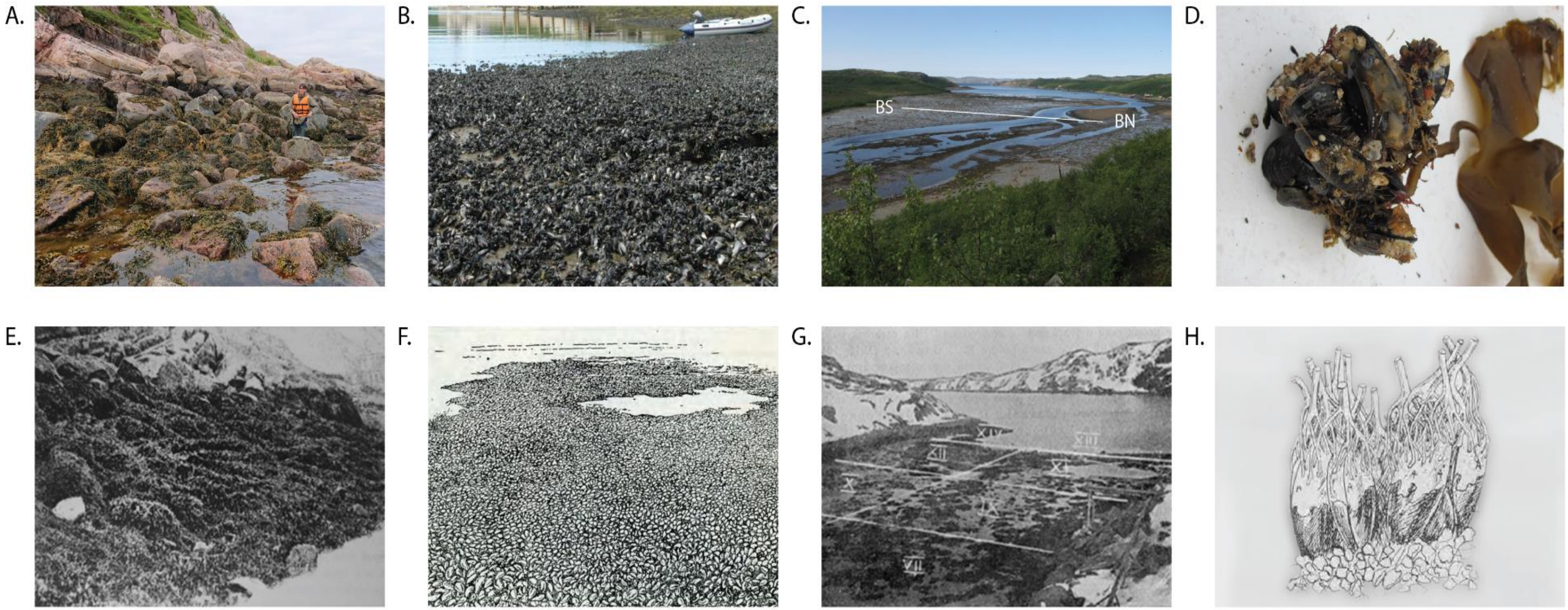


**SFigure 1. Murman mussels and their habitats.** a-d. Photos of characteristic habitats of the Tyuva mussels taken in 2009-2018. (a) Rocky littoral, transect MoS, 2018. (b) Mussel bed, locality SN+0.5, 2009. (c) Littoral sandbanks, location of transect BS is shown. (d) Thallus of sublittoral algae *Alaria esculenta* with attached blue mussels collected at locality MoS-1.5, 2012. e-g. Illustrations of the same habitats in Ekaterininskaya Gavan from the 1920s. (e) Rocky littoral (from Guryanova et al., 1928). (f) Mussel bed* (from Zenkevich, 1963). (g) Littoral sandbanks (from Guryanova et al., 1928). h. Horse mussels *Modiolus modiolus* in kelp rhizoids (from Guryanova, 1924). * *Although the place and the time of the drawing is not specified, it is known that its author, Nikolai Kondakov, visited Ekaterininskaya Gavan in 1928 and 1929*.

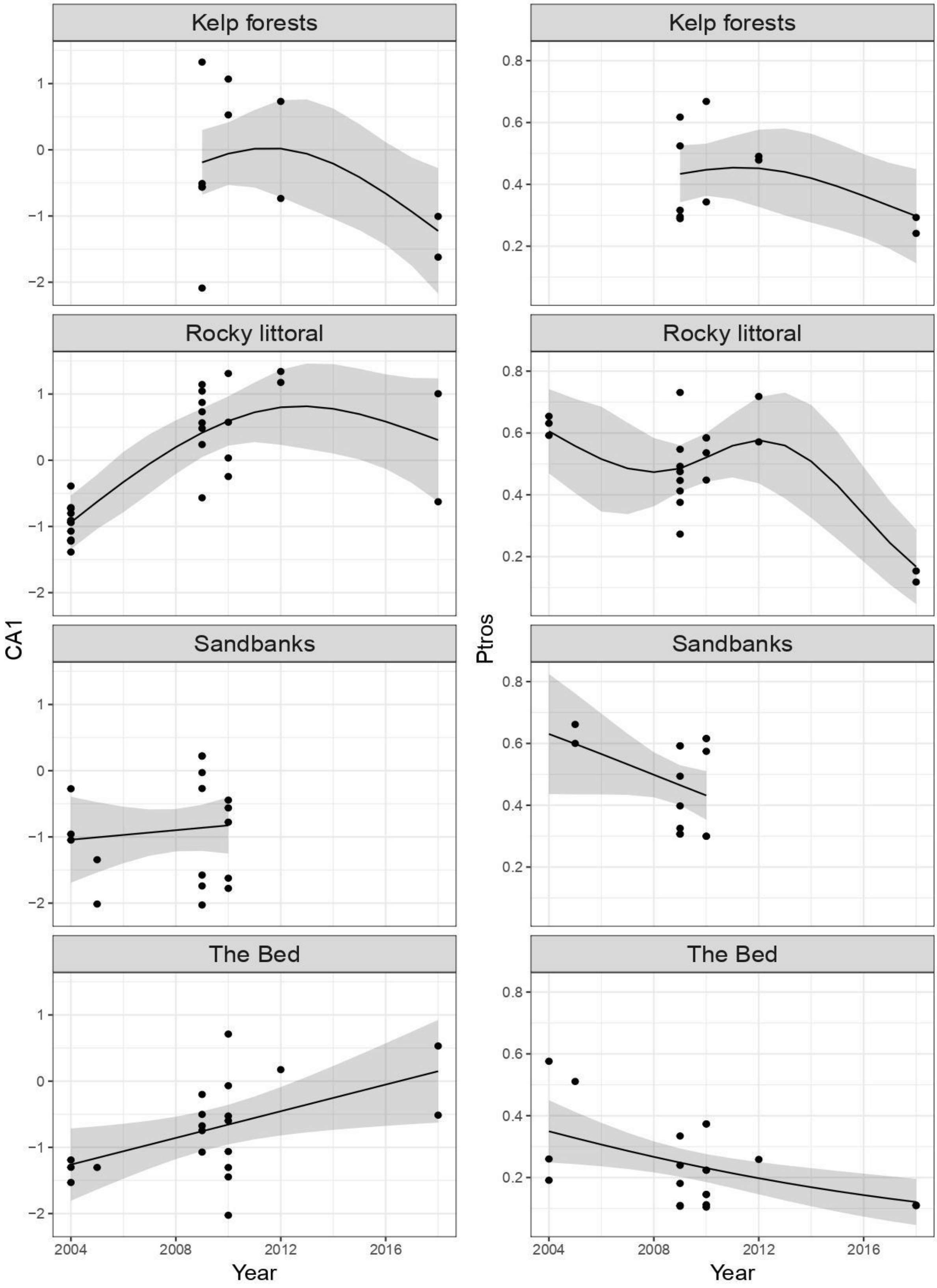


**SFigure 2. Temporal changes in CA1 scores (left) and *Ptros* (right) in different habitats.** Points are empirical values. GAM regression lines with 95% confidence intervals are presented.

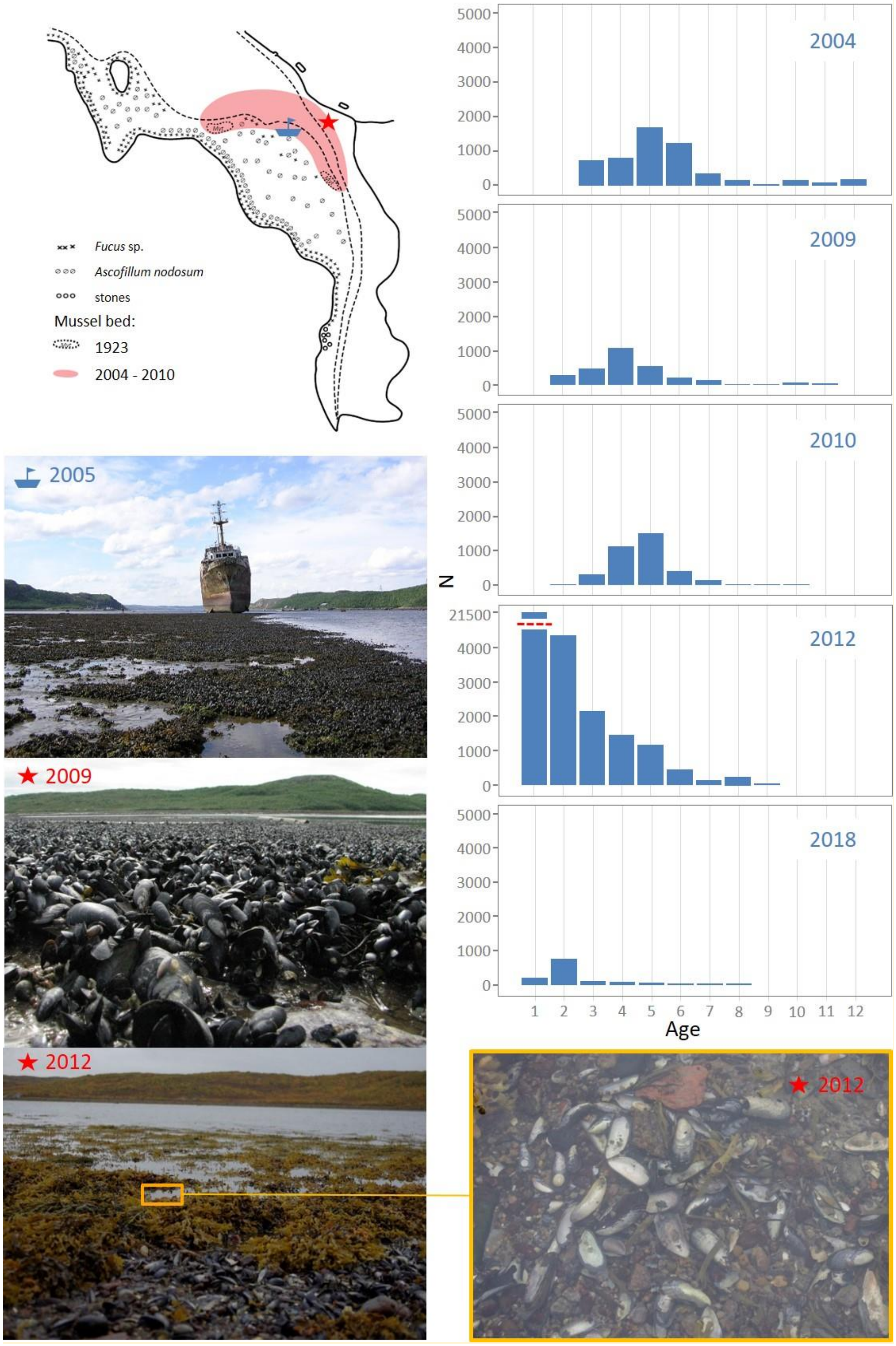


**SFigure 3. Temporal instability of the Bed.** Schematic map of the littoral communities of the top of the Tyuva Inlet in 1923, from Guryanova et al., 1928. The boundaries of "dense mussel settlements" in 1923 are outlined. The map also shows the approximate boundaries of the Bed in 2004-2010 (in red) and the locations seen in the photos. Photographs: selected areas of the Bed in 2009 (near locality BS+05) and in 2012. Bar graphs: age structures of mussels in BS+05 in different years.
